# PILS proteins provide a homeostatic feedback on auxin signaling output

**DOI:** 10.1101/2022.04.28.489893

**Authors:** Elena Feraru, Mugurel I. Feraru, Jeanette Moulinier-Anzola, Maximilian Schwihla, Jonathan Ferreira Da Silva Santos, Lin Sun, Sascha Waidmann, Barbara Korbei, Jürgen Kleine-Vehn

**Author notes:** Correspondence should be addressed to E.F. or J.K.V.

## Abstract

Auxin is a crucial regulator of plant growth and development. Multiple internal and external signals converge at the regulation of auxin metabolism, intercellular transport, and signaling (Pernisova and Vernoux, 2021; Anfang and Shani, 2021). Considering this complexity, it remains largely unknown how plant cells monitor and ensure the homeostasis of auxin responses. PIN-LIKES (PILS) intracellular auxin transport facilitators at the endoplasmic reticulum (ER) are suitable candidates to buffer cellular auxin responses, because they limit nuclear abundance and signaling of auxin (Barbez et al., 2012; Beziat et al., 2017; Feraru et al., 2019; Sun et al., 2020). We used forward genetics to identify mechanisms that define the PILS6 protein abundance and thereby auxin signaling outputs. We screened for *<u>g</u>loomy <u>a</u>nd <u>s</u>hiny <u>p</u>ils* (*gasp*) mutants that define the levels of PILS6-GFP under a constitutive promoter. In this study, we show that *GASP1* encodes for an uncharacterized RING/U-box superfamily protein and impacts on auxin signaling output. We conclude that the low auxin signaling in *gasp1* mutants correlates with reduced abundance of PILS proteins, such as PILS5 and PILS6, which consequently balances auxin-related phenotypes. In agreement, we show that high and low auxin conditions increase and reduce PILS6 protein levels, respectively. Accordingly, non-optimum auxin concentrations are buffered by alterations in PILS6 abundance, consequently leading to homeostatic auxin output regulation. We envision that this feedback mechanism provides robustness to auxin-dependent plant development.

## Results and Discussion

### Forward genetic screen for potential regulators of PILS6

Auxins play a cardinal role in plant growth control. Intercellular auxin transport is crucial for the graded tissue distribution of auxin and thereby provides positional cues. While we have a comprehensive understanding of the tissue distribution of auxin, we still lack a basic understanding of subcellular distribution and signaling of auxin. PIN-LIKES (PILS) are putative intracellular auxin transporters and induce intracellular auxin accumulation at the endoplasmic reticulum (ER) (Barbez et al., 2012). PILS proteins repress the nuclear abundance and signaling of auxin (Barbez et al., 2012; Beziat et al., 2017; Feraru et al., 2019; Sun et al., 2020), presumably by restricting auxin diffusion into the nucleus. Moderately high temperature induces PILS6 protein turnover, which consequently mediates auxin-dependent root thermomorphogenesis (Feraru et al., 2019; Fonseca de Lima et al., 2021), indicating that posttranslational mechanisms define PILS activity and thereby plant adaptation. To further address these uncharted aspects of plant development, we performed a forward genetic screen, using a constitutive PILS6 expression line fused to GFP (*p35*::*PILS6-GFP;* hereafter named *PILS6*^*OE*^), and screened the progeny of about 5,000 M1 ethyl methanesulfonate (EMS)-mutagenized *PILS6*^*OE*^ seeds (Figure 1A). We germinated the *PILS6*^*OE*^ seedlings under standard growth conditions (21 °C) for three days and, subsequently, shifted the plates for 24 h to 29 °C. Then, we evaluated the temperature-sensitive PILS6-GFP fluorescence intensity using an epifluorescence microscope. After re-screening, we identified 21 mutants that showed either reduced (8) or enhanced (13) PILS6-GFP fluorescence intensity under these conditions. We accordingly named these mutants *<u>g</u>loomy <u>a</u>nd <u>s</u>hiny <u>p</u>ils* (*gasp*) (Figure 1A).

**Figure 1.**
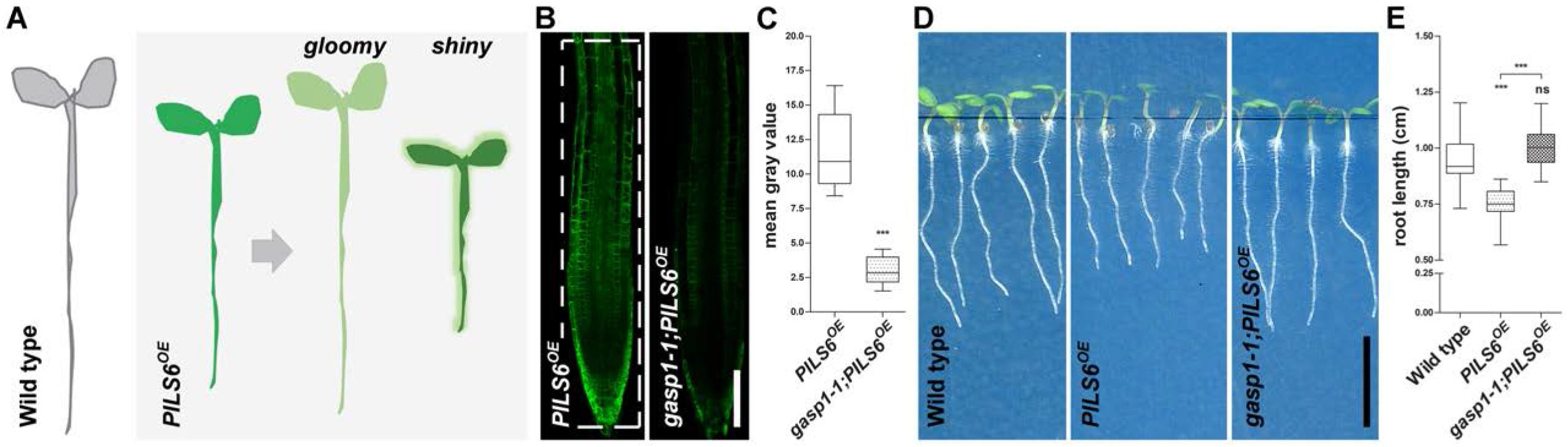
Forward genetic screen for GASP regulators of PILS6 function. A.Graphical representation of the forward genetic screen performed for the identification of *gasp* mutants. EMS-mutagenized seedlings of *PILS6*^*OE*^ were grown for four days under standard growth conditions of 21 °C and subsequently transferred for 24 h to moderately high temperature (29 °C). The seedlings showing either weaker (gloomy) or stronger (shiny) PILS6-GFP signal than *PILS6*^*OE*^ were selected, confirmed, and identified as *gasp* mutants of *PILS6*^*OE*^. Overall, the *gloomy* and *shiny* mutants displayed enhanced and reduced growth when compared with *PILS6*^*OE*^, respectively. B-E.*gasp1-1* mutant affects *PILS6*^*OE*^ fluorescence and general growth under standard conditions of light and temperature. Confocal images (B) and quantification of signal intensity (C) show that *gasp1-1* reduces PILS6-GFP fluorescence. Scans (D) and quantification (E) of root growth at 5 DAG show that *gasp1-1* rescues the short root phenotype of *PILS6*^*OE*^. n = 17, 15 (C) and 19 (E); ns = not significant, ***P < 0.0001, t-test and Mann-Whitney test (C) and One-way ANOVA and Tukey’s multiple comparison test (E). Scale bars, 100 μm (B) and 0,5 cm (D). The white, dashed rectangle shows the ROI used to quantify the signal intensity.

### *gasp1* is a suppressor of PILS6

Among the eight mutants having reduced PILS6-GFP fluorescence signal intensity, we identified the *gasp1-1;PILS6*^*OE*^ mutant that showed almost no PILS6-GFP fluorescence signal after 24 h exposure to 29 °C (Supplemental Figure 1). Notably, when grown under standard temperature of 21 °C, *gasp1-1* mutation caused already a dramatic (85 %) reduction of PILS6-GFP fluorescence intensity when compared to wild type backgrounds (Figures 1B, 1C, Supplemental Figure 1). This finding indicates that the *gasp1-1* mutation affects PILS6-GFP protein abundance independently of moderately high temperature. In accordance with its negative effect on the fluorescence intensity of PILS6-GFP, *gasp1-1* mutation alleviated the short root phenotype of *PILS6*^*OE*^ by 15 % (Figures 1D, 1E). Therefore, we identified *gasp1-1* mutant as a suppressor of PILS6 under standard growth temperature.

### *GASP1* encodes for a RING/U-box superfamily gene

To identify the causal *GASP1* gene, we established a pool of *gasp1-1;PILS6*^*OE*^ individuals isolated from a F2 backcross (*gasp1-1;PILS6*^*OE*^ crossed to *PILS6*^*OE*^). We, accordingly, re-sequenced the genome of this pooled mutant population as well as a pool of non-mutagenized *PILS6*^*OE*^ control seedlings using the Illumina and DNBseq™ platform. By comparing the sequencing results of the two samples, we identified a single nucleotide polymorphism (SNP) in the uncharacterized, protein coding gene AT3G05545, which belongs to the RING/U-box superfamily protein, H2-type (Kraft et al., 2005; Stone et al., 2005). The *gasp1-1* mutation causes a C-to-T mutation, resulting in a proline (P) - to - leucine (L) amino acid substitution at the position 274 (Figure 2A, Supplemental Figure 2A). P274L is not in a conserved region of *GASP1*, but a leucine substitution of a proline residue may have dramatic structural and functional consequences (Vilson et al., 1989; Molnar et al., 2016).

**Figure 2.**
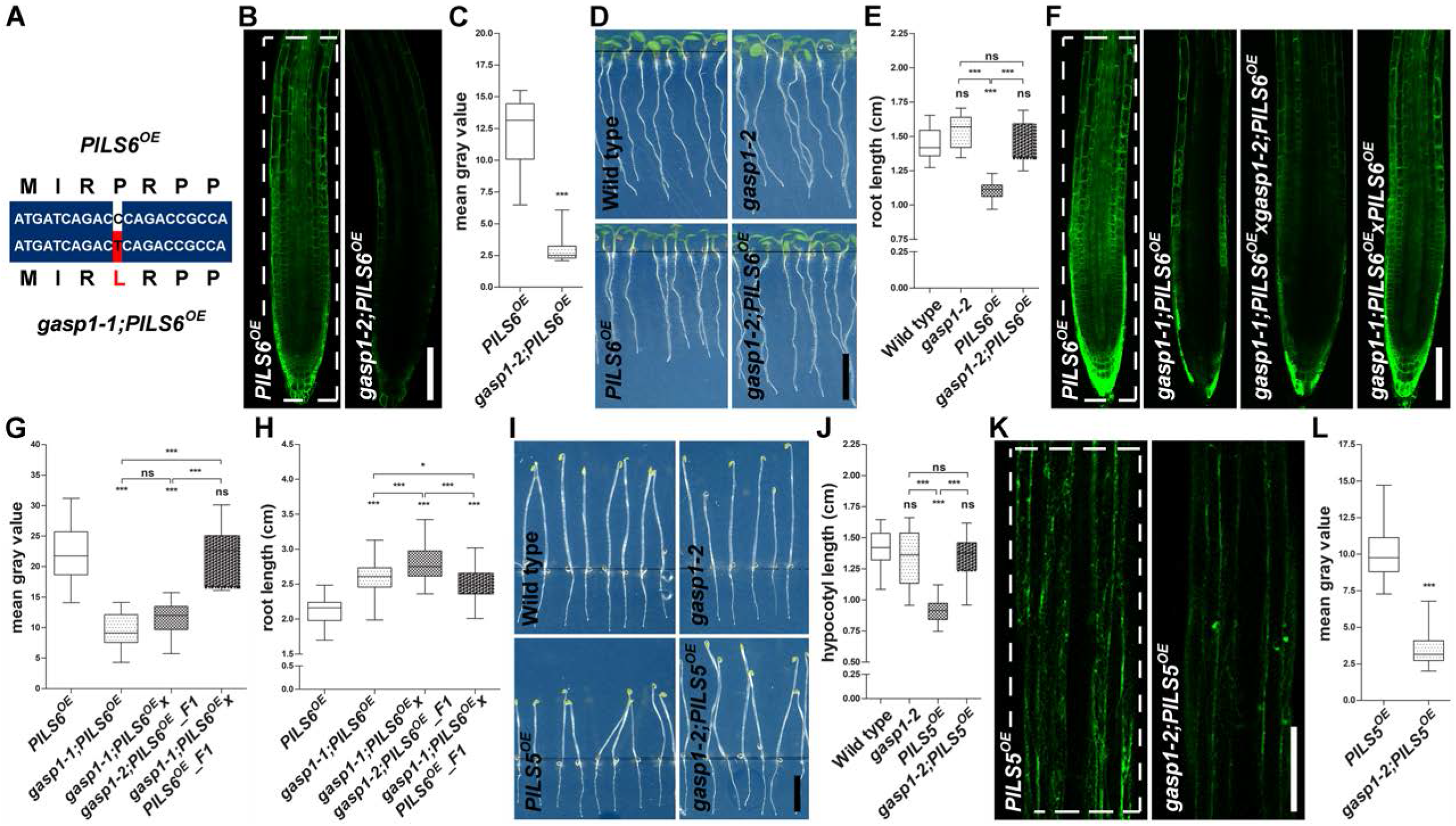
*gasp1* is defective in a RING/U-box superfamily gene. A.Alignment of a short nucleotide sequence from wild-type (top) and mutated (bottom) *GASP* gene. The mutated SNP and the changed amino acid are depicted in red. B-E.The T-DNA insertion *gasp1-2* allele mimics the *gasp1-1* EMS mutant. Confocal images (B), scans of light-grown seedlings at 5 DAG (D) and the respective quantifications (C, E) show that *gasp1-2* allele causes dramatic reduction of PILS6-GFP fluorescence in roots (B, C) and rescues the short root growth (D, E) of *PILS6*^*OE*^. n = 15 (C) and 15-19 (E); ***P < 0.0001, t-test and Mann-Whitney test (C) and ns = not significant, ***P < 0.05, One-way ANOVA and Tukey’s multiple comparison test (E). Scale bars, 100 μm (B) and 0,5 cm (D). F-H.Complementation test showing that *gasp* mutants are allelic. Confocal images (F) and quantifications of signal intensity (G) and root length (H) of the F1 crosses between *gasp1-1* and *gasp1-2* alleles in the *PILS6*^*OE*^ backgrounds and the respective controls show that F1 *gasp1-1;PILS6*^*OE*^x*gasp1-2;PILS6*^*OE*^ causes PILS6-GFP reduction in roots (F, G) and rescues the root growth defects of *PILS6*^*OE*^ (H). n = 10-12 (G) and 67-84 (H); ns = not significant, *P and ***P < 0.05, One-way ANOVA and Tukey’s multiple comparison test (G, H). Scale bar, 100 μm (F). I-L.*gasp1-2* affects *PILS5*^*OE*^ hypocotyl phenotype and PILS5-GFP fluorescence. Scans (I), confocal images (K) and quantifications of hypocotyl length (J) and signal intensity (L) show that *gasp1-2* mutant rescues the phenotype (I, J) and reduces PILS5-GFP signal intensity (K, L) the dark-grown hypocotyls of *PILS5*^*OE*^. n = 20-22 (J) and 15, 16 (L); ns = not significant, ***P < 0.05, One-way ANOVA and Tukey’s multiple comparison test (J) and ***P < 0.0001, t-test and Mann-Whitney test (L). Scale bars, 0.5 cm (I) and 100 μm (K). The white, dashed rectangles show the ROIs used to quantify the signal intensity.

To confirm that *gasp1-1* mutation in AT3G05545 gene is indeed responsible for the suppression of *PILS6*^*OE*^, we isolated a second mutant allele (SALK_091345; hereafter called *gasp1-2*) from the Salk collection of T-DNA insertion lines (Alonso et al. 2003) (Supplemental Figure 2A). *GASP1* transcripts were not detectable in the *gasp1-2* allele, indicating a full knockout of *GASP1* (Supplemental Figure 2B). When crossed to *PILS6*^*OE*^ (*gasp1-2;PILS6*^*OE*^), *gasp1-2* reduced PILS6-GFP fluorescence intensity and, consequently, rescued total root length (Figures 2B-2E). Next, we crossed *gasp1-1;PILS6OE* to the *gasp1-2;PILS6OE* allele as well as to the *PILS6OE* control line. In contrast to the control cross, the allelic test between *gasp1-1* and *gasp1-2* showed that the PILS6-GFP intensity and PILS6^OE^ phenotypes remained suppressed in the F1 generation (Figures 2F-2H; Supplemental Figure 2C). Altogether, we concluded that defects in the *GASP1* are responsible for the phenotypes observed in the *gasp1-1;PILS6OE*.

### *GASP1* defines PILS5 and PILS6 protein abundance

To assess the specificity of the GASP1, we crossed *gasp1-2* to the PILS5 overexpression line (*p35S::PILS5-GFP; PILS5*^*OE*^). Similar to *PILS6*^*OE*^, PILS5-induced reduction in main root growth was also suppressed in *gasp1-2;PILS5*^*OE*^ (Supplemental Figures 2D, 2E). PILS5 overexpression also represses dark-grown hypocotyl growth (Barbez et al., 2012; Beziat et al., 2017), which was as well alleviated by the *gasp1-2* mutation (Figures 2I, 2J). In agreement, the PILS5-GFP signal intensity was strongly reduced in *gasp1-2*;*PILS5OE* dark-grown hypocotyls (Figures 2K, 2L), showing that GASP1 affects at least two PILS proteins, in distinct tissues and growth conditions.

To directly address whether the GASP1 indeed affects PILS5 and PILS6 protein abundance, we subsequently used quantitative western blots. In accordance with the reduced PILS5/6-GFP fluorescence intensity, *gasp1* mutants displayed reduced PILS5 and PILS6 protein levels in the dark-grown hypocotyls and light-grown seedlings, respectively (Figure 3A; Supplemental Figures 3A, 3B). We, accordingly, conclude that GASP1 defines the abundance of PILS proteins, such as PILS5 and PILS6.

**Figure 3.**
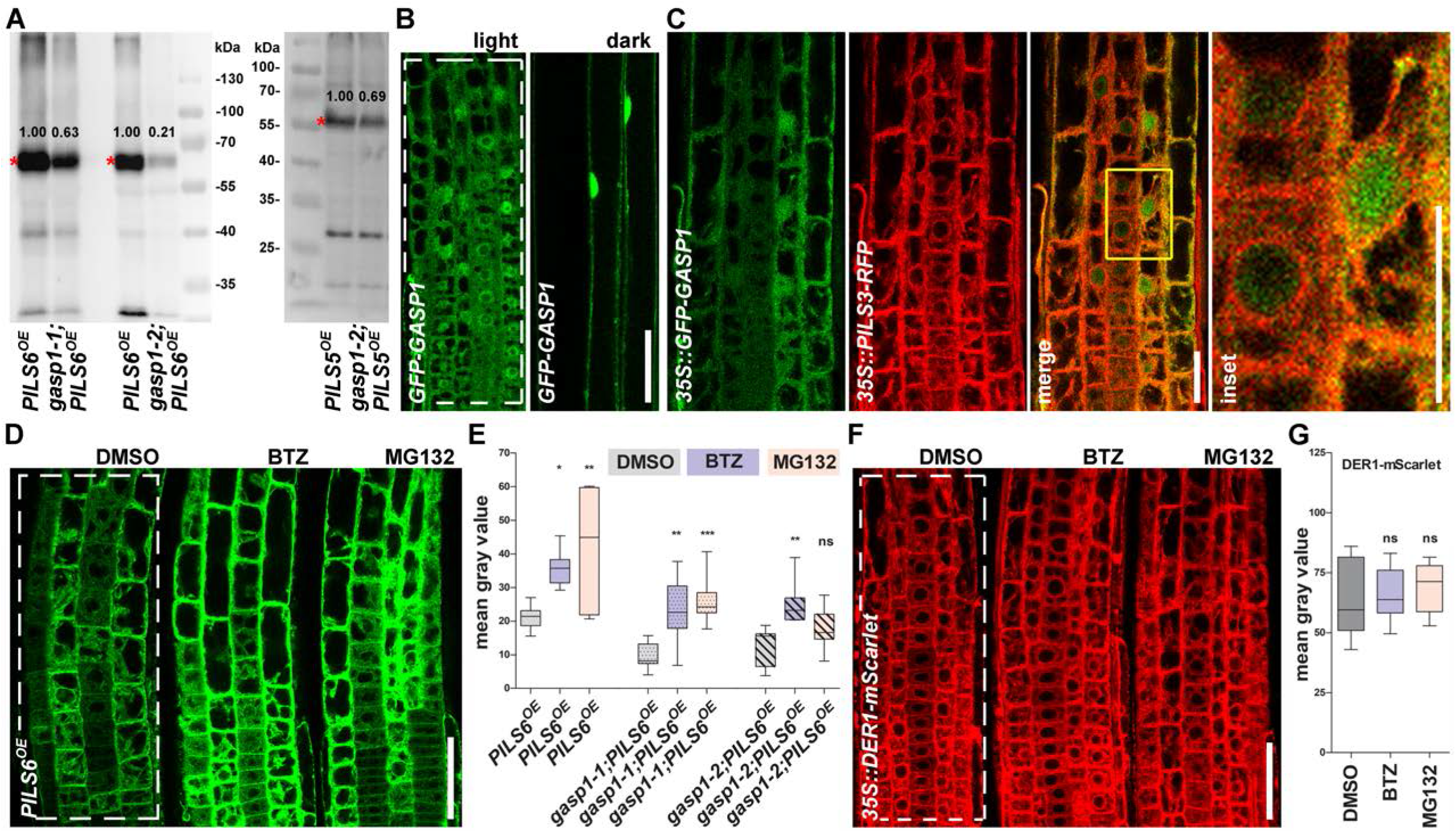
GASP1 is an indirect regulator of PILS5 and PILS6 protein abundance. A.Western blots showing detection of PILS6-GFP (in light-grown seedlings, left image) and PILS5-GFP (in dark-grown hypocotyls, right image). Note the decreased abundance of both PILS5- and PILS6-GFP in the *gasp1* mutants. Red asterisks show PILS5- and PILS6-GFP bands. The values written above the GFP bands represent the intensities that were normalized to the tubulin (left image) or Coomassie (right image) bands presented in Supplemental Figures 3A or 3B, respectively. B,C.*35S::GFP-GASP1* localizes to the cytosol and nucleus. GFP-GASP1 localization is shown in the roots (left) and hypocotyls (right) of light- and dark-grown seedlings from a homozygous, F3 generation, respectively (B). GFP-GASP1 does not colocalizes with the ER marker PILS3-RFP in a F1 cross (C). The yellow rectangle shows the region that is magnified in the inset. Scale bars, 50 μm (B) and 25 μm (C). D,E.Proteasome inhibitors stabilize PILS6-GFP independently of GASP1. Confocal images (D) and quantification of signal intensity (E) show that a short treatment (3 h) with the proteasome inhibitors BTZ [50 uM]) or MG132 [50 uM] stabilizes PILS6-GFP in WT (D, E) and in *gasp1* mutants (E). n = 7-9; ns = not significant, *P, **P, ***P < 0.05, One-way ANOVA and Tukey’s multiple comparison test (E). Scale bar, 50 μm (D). F,G.Proteasome inhibitors do not affect DER1-mScarlet. Confocal images (F) and quantification of signal intensity (G) show that a 3 h treatment with the proteasome inhibitors BTZ [50 uM] or MG132 [50 uM] does not affect DER1-mScarlet fluorescence intensity. n = 11-13; ns = not significant, One-way ANOVA and Tukey’s multiple comparison test (G). Scale bar, 50 μm (F). The white, dashed rectangles show the ROIs used to quantify the signal intensity.

*GASP1* belongs to the RING/U-box superfamily and plays a role as an E3 ubiquitin ligase (Kraft et al., 2005; Stone et al., 2005). RING E3 ubiquitin ligases typically mediate the ubiquitination of target proteins, where K48-linked ubiquitination recruits these targets for degradation via the 26S proteasome (Joazeiro and Weissman, 2000; Smalle and Vierstra, 2004; Vierstra, 2009). To address if GASP1 could directly modulate PILS proteins abundance at the ER membrane, we generated a transgenic line overexpressing GFP-GASP1 fusion. *35S::GFP-GASP1* lines displayed weak but ubiquitous signal in the root (Supplemental Figure 3C). In agreement with its predicted localization (https://suba.live/suba-app/factsheet.html?id=AT3G05545; Hooper et al., 2017), GFP-GASP1 was detectable in the nucleus, but showed also cytosolic localization (Figure 3B). Although the GFP-GASP1 appeared enriched in the perinuclear regions of light-grown seedlings, we did not detect pronounced association with the ER and, accordingly, GFP-GASP1 did not show co-localization with PILS3-RFP (Figures 3B, 3C). In addition, GASP1 did not interact with PILS3 or PILS5 proteins in a yeast mating-based split-ubiquitin system (Supplemental Figure 3D). Even though we cannot fully rule out a direct interaction *in planta*, we assume that the putative E3 ubiquitin ligase GASP1 rather indirectly affects the protein abundance of PILS5 and PILS6. Considering that E3 ligases are typically negative regulators of their clients, the reduced PILS5 and PILS6 abundance in *gasp1* mutants also questions the direct impact of the E3 ubiquitin ligase GASP1. Either GASP1 plays an unusual role for an E3 ligase or it defines the ubiquitination and, hence, degradation of cytosolic and/or nuclear proteins that are upstream regulators of PILS5 and PILS6 via the ubiquitin-26S proteasome pathway. To test if the degradation of the PILS proteins is affected by the disturbance of the ubiquitin-26S proteasome pathway, we subsequently used MG132 and Bortezomib (BTZ) to pharmacologically interfere with the proteasome function in *Arabidopsis* (Marshall et al., 2015; Yu et al., 2015; Wang et al., 2016). Seedlings of *PILS6OE* were treated for three hours with MG132 or BTZ, which caused a significant increase of PILS6-GFP fluorescence in roots when compared to the DMSO-treated control seedlings (Figure 3D, 3E), indicating that PILS6 protein abundance is regulated in a 26S proteasome-sensitive manner. Importantly, another ER membrane-localized protein, a component of the ERAD pathway, the RING E3 ligase DERLIN1 (Kirst et al., 2005) translationally fused to mSCARLET (*35S::DER1-mSCARLET*), is not affected by proteasome inhibition, indicating some specificity of this effect (Figures 3F, 3G). The pharmacological inhibition of the 26S proteasome also increased the PILS6-GFP fluorescence in *gasp1-1* and *gasp1-2* mutants (Figure 3E; Supplemental Figures 3E-3G), indicating that other molecular components contribute to the proteasome effect on PILS6 abundance. Collectively, our data shows that although the ubiquitin-26S proteasome activity is required for PILS6 degradation, PILS protein abundance is not directly controlled by the E3 ligase GASP1.

### Auxin signaling modulates PILS6 protein abundance

We next addressed whether the reduced PILS5 and PILS6 protein abundance correlates with the expected increased nuclear auxin signaling output in *gasp1* (Barbez et al., 2012; Beziat et al., 2017; Feraru et al., 2019; Sun et al., 2020). To visualize the auxin signaling output in *gasp1-2*, we introduced the auxin response marker *DR5::GFP* by crossing. While *DR5::GFP* signal intensity was not distinguishable in the root tip of *gasp1-2* mutant and wild type (Supplemental Figures 4A, 4B), we, unexpectedly, observed reduced *DR5::GFP* signal in the upper vascular tissues of light-grown roots as well as dark-grown hypocotyls of *gasp1-2* mutant (Figures 4A-4D). In agreement, auxin responsive genes, such as *IAA1, IAA5, IAA7, SAUR19*, and *SAUR63*, showed reduced expression in the light-grown seedlings and in the dark-grown hypocotyls of *gasp1* mutants (Figures 4E, 4F). This finding suggests that *GASP1* is required to maintain auxin signaling output in roots and shoots, which is independent from its effect on PILS5 and PILS6 abundance (Figures 4E, 4F). The overexpression of PILS proteins also limits nuclear auxin signaling, but the repression of auxin signaling output was not additive in *gasp1-1;PILS6*^*OE*^, *gasp1-2;PILS6*^*OE*^, and *gasp1-2;PILS5*^*OE*^ (Figures 4E, 4F), suggesting that the effect on PILS abundance balances the auxin response. This finding hints at a molecular mechanism in which the PILS abundance could relate to a homeostatic feedback mechanism on auxin signaling output.

**Figure 4.**
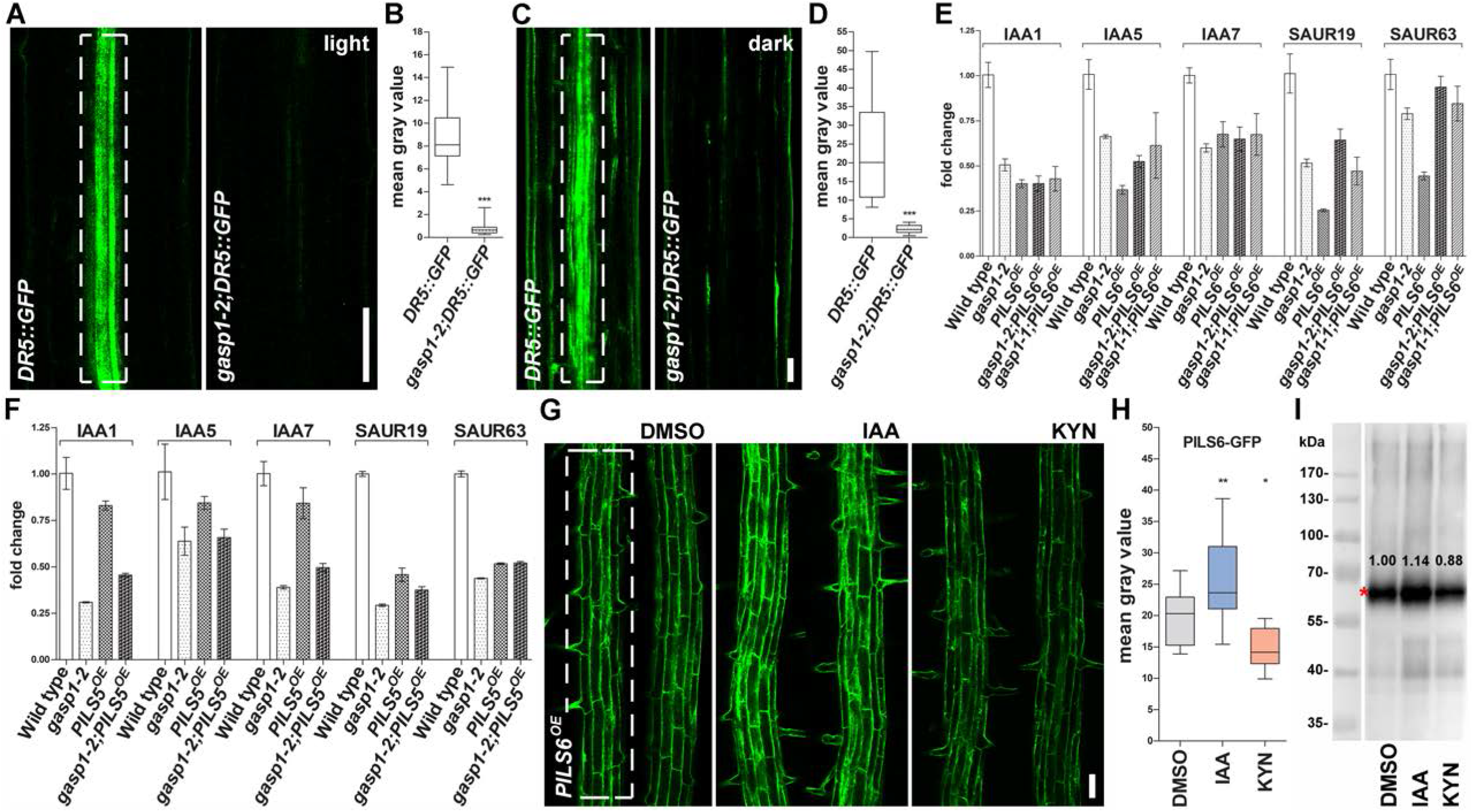
Auxin signaling affects PILS6 protein abundance. A-F.Auxin signaling is reduced in the *gasp1* mutants. Confocal images (A, C) and quantifications of signal intensity (B, D) show that *gasp1-2* mutation negatively affects *DR5::GFP* signal in the roots (differentiation zone is presented) (A, B) of light- and hypocotyls (C, D) of dark-grown seedlings. qPCR analysis showing the expression of some IAA and SAUR genes in the entire seedlings grown in the light (E) and hypocotyls of seedlings grown in the dark (F). Note the reduced expression of the auxin responsive genes in all transgenic lines. n = 16, 17 (B) and 9, 10 (D); ***P < 0.0001, t-test and Mann-Whitney test (B, D). Scale bars, 50 μm (A, C). G-I.Auxin signaling modulates PILS6 protein abundance. Confocal images (G), quantification of signal intensity (H) and immunoblot with anti-GFP (I) show that 24 h treatment with either [100 nM] IAA or [1uM] KYN increases or reduces PILS6-GFP abundance in roots of light-grown *PILS6*^*OE*^ seedlings. Red asterisk (I) marks PILS6-GFP bands. The values written above the GFP bands (I) represent the intensities that were normalized to the tubulin bands presented in Supplemental Figure 4C. n = 17; *P, **P < 0.05, One-way ANOVA and Tukey’s multiple comparison test (H). Scale bar, 50 μm (G). The white, dashed rectangles show the ROIs used to quantify the signal intensity.

This prompted us to address whether the diminished auxin signaling output observed in *gasp1* mutants could reflect an auxin impact on PILS proteins abundance. To test this, we used L-Kynurenin (KYN) to pharmacologically interfere with auxin biosynthesis and hence signaling (He at al., 2011). KYN applications indeed phenocopied *gasp1* mutants and decreased the PILS6-GFP fluorescence as well as protein abundance (Figures 4G-4I; Supplemental Figure 4C). Conversely, 100 nM IAA treatment increased both PILS6-GFP fluorescence and protein abundance (Figures 4G-4I; Supplemental Figure 4C). These experiments suggest that high and low auxin signaling outputs in- and de-crease PILS6 abundance, respectively. In contrast, the ER membrane marker DER1-mScarlet did not show any response to the treatments with either KYN or IAA (Supplemental Figures 4D, 4E), suggesting certain specificity of this auxin response. This data proposes that auxin exerts a homeostatic feedback on its own signaling rate by controlling the abundance of PILS intracellular auxin transporters.

### Concluding remarks

Our forward genetic screen performed to identify regulators of PILS6 protein turnover under moderately high temperature yielded 21 *gasp* mutants that either decreased (*gloomy*) or increased (*shiny*) *PILS6*^*OE*^ traits. In this study, we investigated *gasp1-1* that repressed PILS6-GFP already under standard temperature and found that *GASP1* may function as a modulator of auxin signaling rates. *GASP1* encodes for an uncharacterized E3 ubiquitin ligase, from the H-type, that belongs to the RING/U-box superfamily protein, which supposedly mediates substrate specific ubiquitination (Kraft et al., 2005; Stone et al., 2005). The *gasp1* mutants display severe reduction in auxin signaling output, but in contrast merely increase phenotypic trait variations, and are largely not distinguishable from wild type seedlings. It is hence conceivable that homeostatic auxin responses may balance the molecular responses in *gasp1*. The biological role of *GASP1* remains largely unknown, but we show here that the severely reduced auxin signaling output in *gasp1* mutants is in part compensated by enhanced turnover of at least PILS5 and PILS6 proteins. Our data shows that GASP1 does not directly interact with PILS proteins, such as PILS3 or PILS5 heterologously expressed in yeast. We propose that the GASP1 impact on auxin signaling output indirectly affects PILS5 and PILS6 turnover. We, accordingly, show that sub- and supra-optimum levels of auxin de- and increase PILS6 abundance at the ER, respectively. It remains to be seen how precisely auxin levels determine the turnover of PILS proteins. Such a response could involve the canonical TIR1/AFB auxin receptors and downstream signaling events, altering the yet to be defined PILS degradation mechanisms. Alternatively, auxin availability may structurally affect PILS proteins, which could alter their interaction with the degradation machinery.

PILS proteins define the nuclear abundance and signaling of auxin, which seems highly responsive to internal and external signal perturbations (Beziat et al., 2017; Feraru et al., 2019; Sun et al., 2020). Here, we propose a working model where an auxin impact on PILS abundance provides homeostatic feedback (Supplemental Figure F), enabling auxin signaling output to maintain its own cellular homeostasis. Altogether, we envision that a PILS-dependent feedback mechanism provides robustness to plant growth and development.

## Author contribution

E.F., M.I.F., and J.K.-V. conceived the manuscript. E.F., M.I.F., J.M.-A., M.S., J.F.D.S.S., L.S., and S.W. performed experiments and analyzed data. M.I.F. made the figures. E.F and J.K.-V wrote the manuscript. E.F., B.K., and J.K.-V edited the manuscript.

## Acknowledgements and Funding

We are grateful to BOKU-VIBT Imaging Center and M. Debreczeny for access and advice; L. Mach and R. Strasser for sharing equipment. This work was supported by the Austrian Science Fund (FWF) (Projects P 26591 and P 29754 to J.K.-V., P 30850-B32 to B.K., P 33497 to S.W., and Hertha Firnberg T728-B16 and Elise Richter V690-B25 to E.F.), European Research Council (AuxinER - ERC starting grant to J.K.-V.), and the German Science Fund (DFG) (Project 470007283 and CIBSS – EXC-2189 – Project ID 390939984 to J.K.-V.).

## Material and methods Plant material

*Arabidopis thaliana* ecotype Col-0 (wild-type), *p35S::PILS5-GFP* (*PILS5*^*OE*^; Barbez et al., 2012), *p35S::PILS6-GFP* (*PILS6*^*OE*^; Barbez et al., 2012), *p35S::PILS3-RFP* (Barbez et al., 2012 and this study), *pDR5rev::GFP* (Benkova et al., 2003) were previously described. *gasp1-2* (SALK_091345) was obtained from NASC (Alonso et al., 2003); *gasp1-1* was identified in this study.

### Growth conditions

70 % ethanol-sterilized (1-2 min sterilization in paper bags, followed by 30 minutes drying) seeds were plated usually on one single line, uniformly spaced, in the upper part of Petri dishes containing 50 ml solidified Murashige and Skoog (MS) agar medium (0.8 % agar, 0.5 x MS, and 1 % sucrose, pH 5.9), then stratified for 3 days in the dark at 4 °C, and grown on vertically oriented plates in a plant cabinet equipped with above placed cool-white fluorescent bulbs and set at about 140 μmol/m^-2^s^-1^, long day photoperiod, and 21 °C. This ensures that all seedlings in that plate are exposed to the same light intensity and humidity, which results in low variability. For HT-related experiments, we used the growth conditions described in Feraru et al. 2019. Seedlings were grown on plates (in pairs), for four days, under 21 °C (standard temperature) and subsequently shifted for 24 h in a cabinet displaying similar settings, excepting the temperature that was 29 °C (moderately high temperature). The control plates remained in the cabinet equipped with standard conditions.

### EMS mutagenesis, forward genetic screen, and sequencing

Roughly 10,000 seeds of *35S::PILS6-GFP (PILS6*^*OE*^) were soaked (gently shaking) for 10 hours in 0.1 M phosphate buffer, pH 7, containing 0.3 % (v/v) ethyl methanesulfonate (EMS). Prior mutagenesis, the seeds were soaked for 5 minutes in water containing 0.05 % Triton X, then rinsed three times with water. After mutagenesis, the seeds were rinsed 7 times with water, then dispersed as desired in 0.1 % agarose, and transferred to soil by pipetting. From about 5,400 mutagenized plants (M1), we harvested 360 pools (each pool containing about 15 M1), and screened under an Olympus stereomicroscope over 80,000 M2 seedlings for individual seedlings with weaker or stronger fluorescence than *PILS6*^*OE*^ control. The seedlings having different fluorescence intensity than the control were picked up, propagated, and confirmed in the next generation as *<u>g</u>loomy <u>a</u>nd <u>s</u>hiny <u>p</u>ils* (*gasp*) mutants.

For sequencing of *gasp1-1*, we crossed *gasp1-1;PILS6*^*OE*^ to *PILS6*^*OE*^ and selected in F2 the individuals showing *gasp1-1;PILS6*^*OE*^ phenotype. A pool of seedlings weighting 100 mg was used to extract genomic DNA by using the DNeasy Plant Mini Kit (Qiagen), according to the manufacturer’s instructions. A sample of more than 1.5 ug (> 13 ng/ul sample concentration) was sent for sequencing. Along, we sent a similar sample containing the *PILS6*^*OE*^ control. The samples were sequenced using the DNBseq platform and the standard bioinformatics analysis was performed by the company. The company identified the different SNPs between each sample and *Arabidopsis thaliana* genome from the TAIR database, followed by SNP calling, annotation, and statistics. To identify the *gasp1-1* mutation, we compared the list of SNPs identified in the *gasp1-1;PILS6*^*OE*^ with the list of SNPs identified in the *PILS6*^*OE*^ sample. After elimination of common SNPs between the two samples and of those heterozygous and synonymous SNPs, we identified one single, typical EMS mutation.

### Quantification of phenotypes

For root and hypocotyl length measurements, seedlings were grown on vertically oriented plates in the light (root) or dark (hypocotyl). Plates were scanned with Epson Perfection V700 scanner and the length was measured by using ImageJ 1.41 software (http://rsb.info.nih.gov/ij/).

### Confocal imaging and quantification

Leica TCS SP5 confocal microscope was used for fluorescence imaging. Unless stated differently, 5-day-old seedlings were used. When treated prior imaging, the seedlings were either submerged in MS liquid medium (MG132, BTZ) or transferred on plates (IAA, KYN) containing the desired concentration of the drug or similar amount of solvent and kept in the plant cabinet for the duration specified in the text or figure legend. The mean gray value of the fluorescence intensity was quantified in a defined rectangle region of interest (ROI), marked on the images, by using “Quantify” tool of Leica software (LAS AF Lite).

## Cloning

To generate *p35S::GFP-GASP1*, the GASP genomic fragment was cloned into the pDONR221 by using the primers B1_GASP_FP and B2_GASP^STOP^_RP listed in Table 1. The resulting entry clone was subsequently transferred to the gateway-compatible destination vector pK7WGF2 (Karimi et al., 2002). Transformed lines were selected on 100 mg/L Kanamycin in F1 and 25 mg/L Kanamycin in F2. For the split ubiquitin assay, we amplified the PILS3, PILS5, and GASP1 coding sequence without stop codon by using PILS3_FP, PILS3^NOSTOP^_RP, PILS5_FP and PILS5^NOSTOP^_RP, B1_GASP_FP, and B2_GASP^NOSTOP^_RP listed in Table 1. The fragments were firstly cloned into the pDONR221. Subsequently, we recombined the baits (PILS3 and PILS5) and prey (GASP1) into pMetYC-DEST (Grefen et al. 2007) and PNX35-DEST (Grefen & Blatt, 2012), respectively. We used Gibson Assembly (NEB) to generate *35S::DER1-mSCARLET*. The coding sequence of DER1 (DER1_FP and DER1_RP), the 35S promoter (35S_FP and 35S_RP), and mScarlet-i tag (mScarlet_FP and mScarlet_RP) were amplified by PCR using Q5 High-Fidelity DNA Polymerase (NEB). The fragments were then cloned into a linearized (EcoRV-HF-NEB) pPLV03 vector by using Gibson Assembly. The transformed lines were selected on 15 mg/L Phosphinothricin (Basta). *35S::PILS3-RFP* plasmid generated previously (Barbez et al., 2012) was transformed into Col-0 plants and the transformed lines were selected on 20 mg/L hygromycin.

**Table 1.**
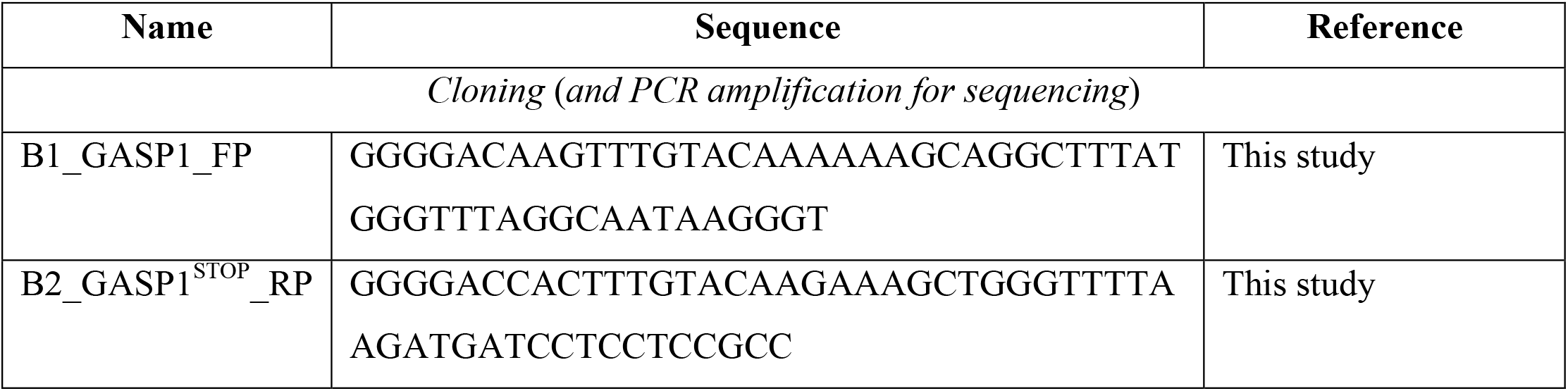
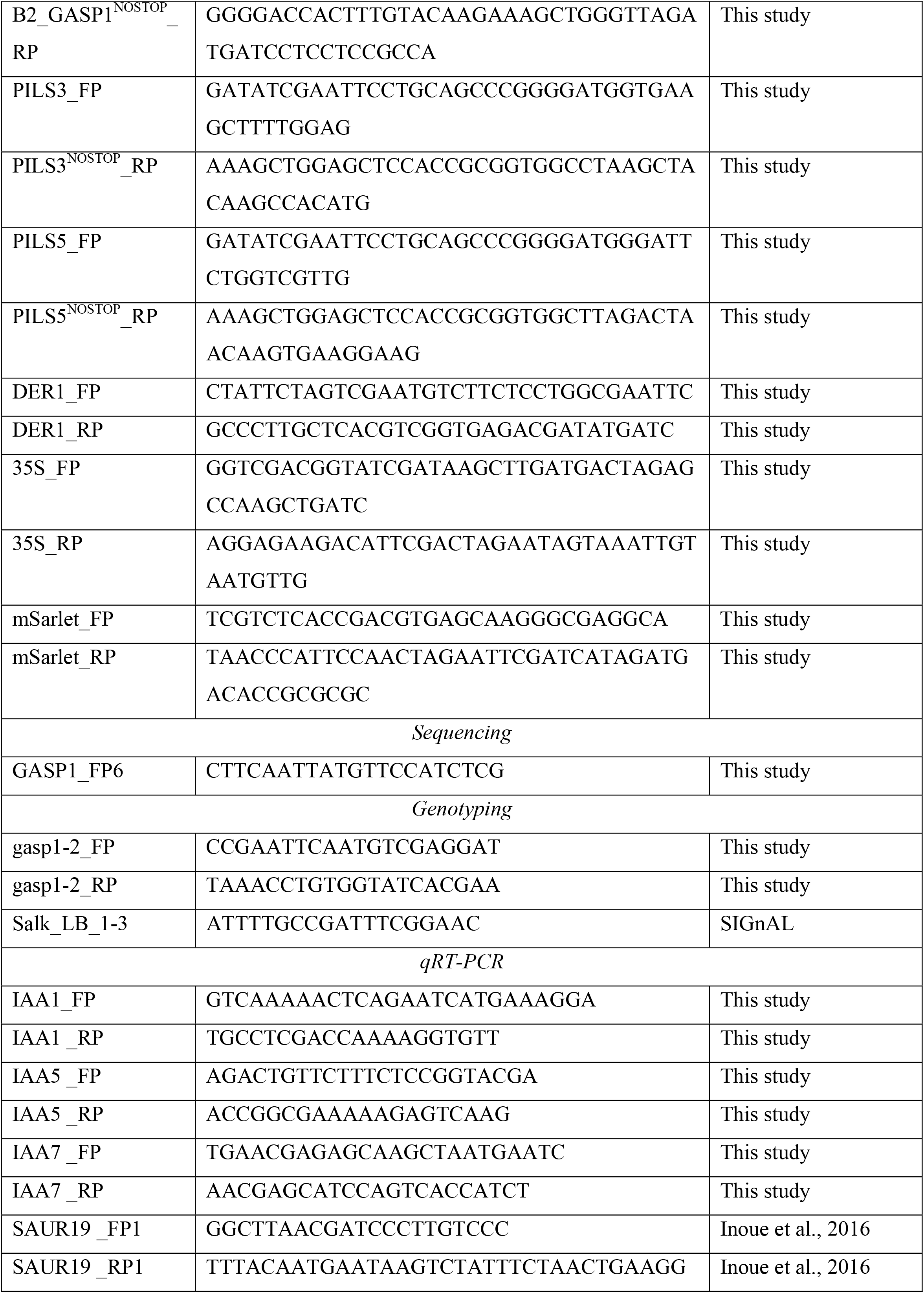
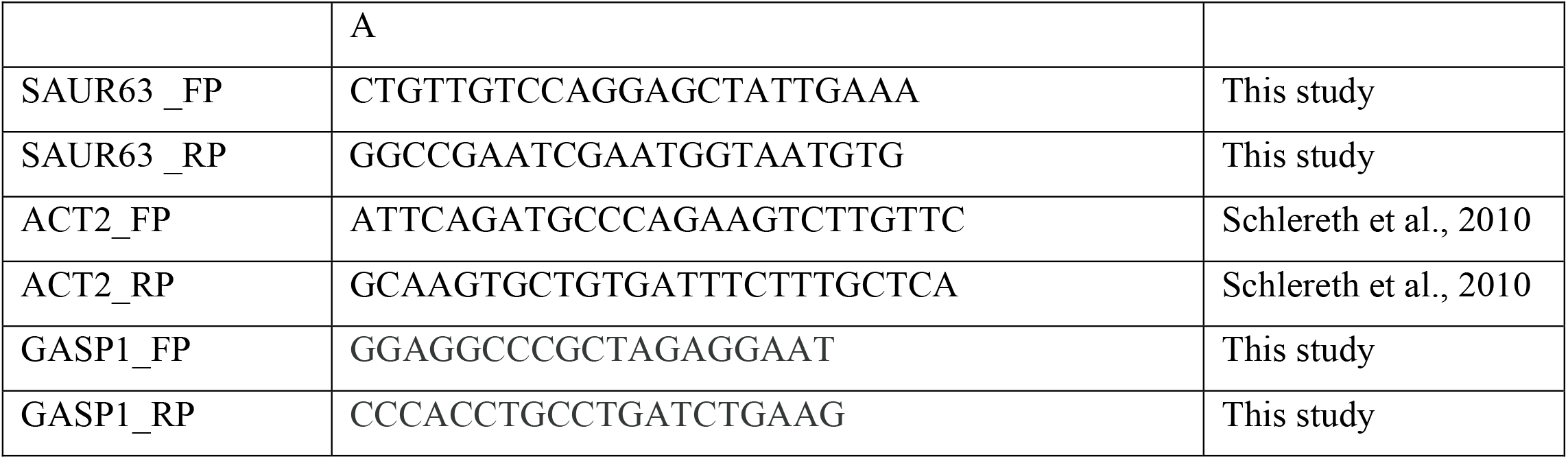
Primers used in this study.

### Split Ubiquitin

For the split ubiquitin assay, the yeast strains THY.AP4 and THY.AP5 were transformed with bait and prey constructs, respectively, using a modified protocol from (Gietz & Woods, 2002). Approximately 100 µl of fresh yeast were scraped from YPD plates and resuspended in 200 µl sterile H_2_O. The resuspended yeast was then centrifuged for 5 minutes at 2000 g and the supernatant removed. The yeast was then resuspended in 200 µl Yeast transformation buffer (40 % PEG 3350, 200 mM LiAc, 100 mM DTT), added 10 µl single stranded carrier DNA and 1 µg of plasmid DNA and mixed by pipetting up and down. We incubated the yeast for 15 min at 30 °C and for 45 min at 45 °C, subsequently plated on SD medium, and incubated for 4 days at 28 °C. A pool of transformed colonies was mated as described in Grefen et al. 2007. The selected diploid colonies were then incubated on plates contacting selective medium (SD -Trp, - Leu, -Ade, -His, -Ura) at 21 °C under light and dark conditions. Growth was recorded up to 9 days after plating.

### Sequencing and genotyping

To identify *gasp1-1*, we amplified the genomic sequence with B1_GASP1_FP and B2_GASP1^STOP^_RP and sequenced the sequence around the mutation by using the primer GASP1_FP6 listed in Table 1. To genotype *gasp1-2*, we used a combination of *gasp1*-*2* and t-DNA insertion-specific primers listed in Table 1.

### qRT-PCR Analysis

It has been performed as described in Feraru et al., 2019. We used the InnuPREP Plant RNA Kit (Analytic Jena) to extract total RNA according to the manufacturer’s recommendation. The RNA samples were treated with InnuPREP DNase I (Analytic Jena). To synthesize cDNA, 1 μg of RNA and the iSCRIPT cDNA Synthesis Kit (Bio-Rad) were used. qRT-PCRs ware carried out in a C1000 Touch Thermal Cycler equipped with the CFX96 Touch Real-Time PCR Detection System (Bio-Rad) and using the Takyon qPCR Kit for SYBER assay (Eurogentec), according to the manufacturer’s recommendation. Gene expression was normalized to the expression of *ACTIN2*. The primers used for qPCRs are listed in Table 1.

### Western Blots

5-day-old dark-grown hypocotyls were used for the experiment related to *PILS5*^*OE*^ and 6-day-old total seedlings for the experiments related to *PILS6*^*OE*^. For IAA and KYN treatments, the seedlings were grown for five days on nylon mesh on MS plates, then transferred with the underlying mesh to the plates supplemented with IAA, KYN or similar amount of DMSO solvent, and harvested after 24 h. Protein extraction was performed as described in Moulinier-Anzola et al., 2020. Each sample contained 20 mg seedlings. The frozen plant material was ground using a Retsch mill and extracted in 150 μl buffer (65 mM Tris [pH 6.8], 8 M urea, 10 % glycerin, 2 % SDS, 5 % β-mercaptoethanol and 0.25 % bromophenol blue). The samples were furthermore heated at 65 °C for 5 min and spun down before loading. Anti-GFP (Roche, 1:1000), monoclonal anti-α-Tubulin (Sigma, 1:100000), and goat anti-mouse IgG (Jackson ImmunoResearch, 1:40000) antibodies were used for detection of *PILS5*^*OE*^, *PILS6*^*OE*^ and Tubulin.

The experiments presented in this study have been performed at least three times or in three replicates.

## Figure Legends

**Figure 1S.**
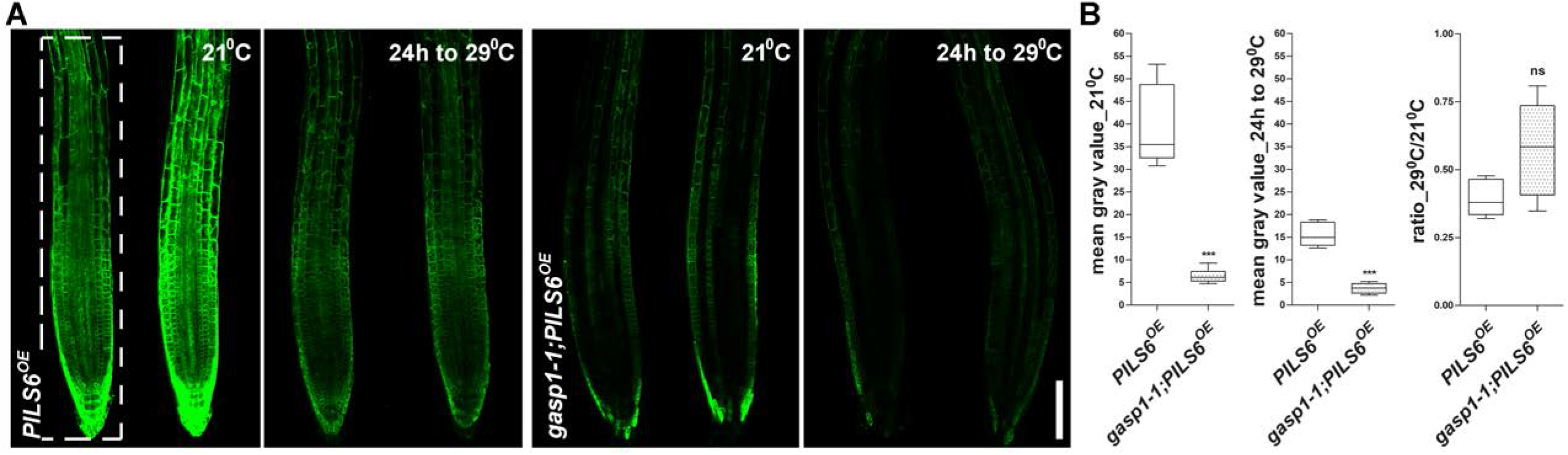
*gasp1-1* regulates PILS6 independently of moderately high temperature. A,B.*gasp1-1* mutation affects *PILS6*^*OE*^ already under standard growth conditions. Confocal images (A) and quantification of signal intensity (B) show that PILS6-GFP fluorescence is already weaker in the *PILS6*^*OE*^ seedlings grown under 21 °C and is further reduced, similarly to the control seedlings, after 24 h exposure to 29 °C. n = 8; ns = not significant, ***P = 0.0007, t-test and Mann-Whitney test (B). Scale bar, 100 μm (A). The white, dashed rectangle shows the ROI used to quantify the signal intensity.

**Figure 2S.**
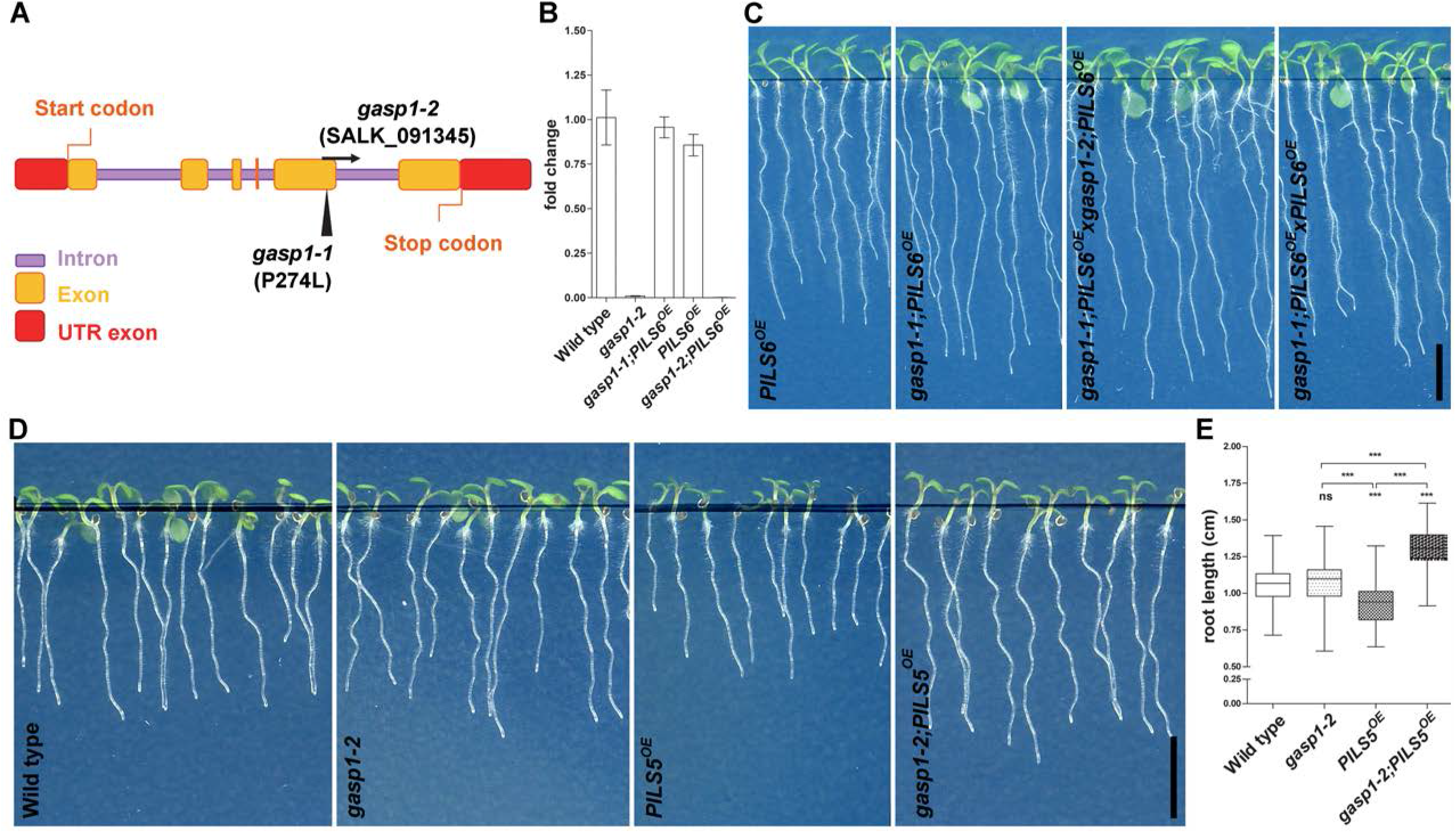
*GASP1* encodes for a RING/U-box superfamily gene. A.Schematic representation of *GASP1* gene, according to PLAZA 5.0 (https://bioinformatics.psb.ugent.be/plaza/versions/plaza_v5_dicots/). Black arrowhead and arrow show the approximate positions of *gasp1-1* SNP and *gasp1-2* t-DNA insertion, respectively. B. qPCR showing *GASP1* transcript. *GASP1* transcript is absent in *gasp1-2* mutant and unchanged in *gasp1-1* mutant. Wild type and *PILS6*^*OE*^ were used as controls. C.*gasp1* mutants are allelic. Scans of 7 DAG seedlings show that the F1 cross between *gasp1-1* and *gasp1-2* mutants in *PILS6*^*OE*^ background rescues the short root growth of *PILS6*^*OE*^. Scale bar, 0.5 cm. D,E.*gasp1-2* affects *PILS5*^*OE*^ root phenotype. Scans (D) and quantification (E) show that *gasp1-2* allele rescues root growth of 5 DAG light-grown *PILS5*^*OE*^ seedlings. n = 41-43; ns = not significant, ***P < 0.05, One-way ANOVA and Tukey’s multiple comparison test (E). Scale bar, 0.5 cm (D).

**Figure 3S.**
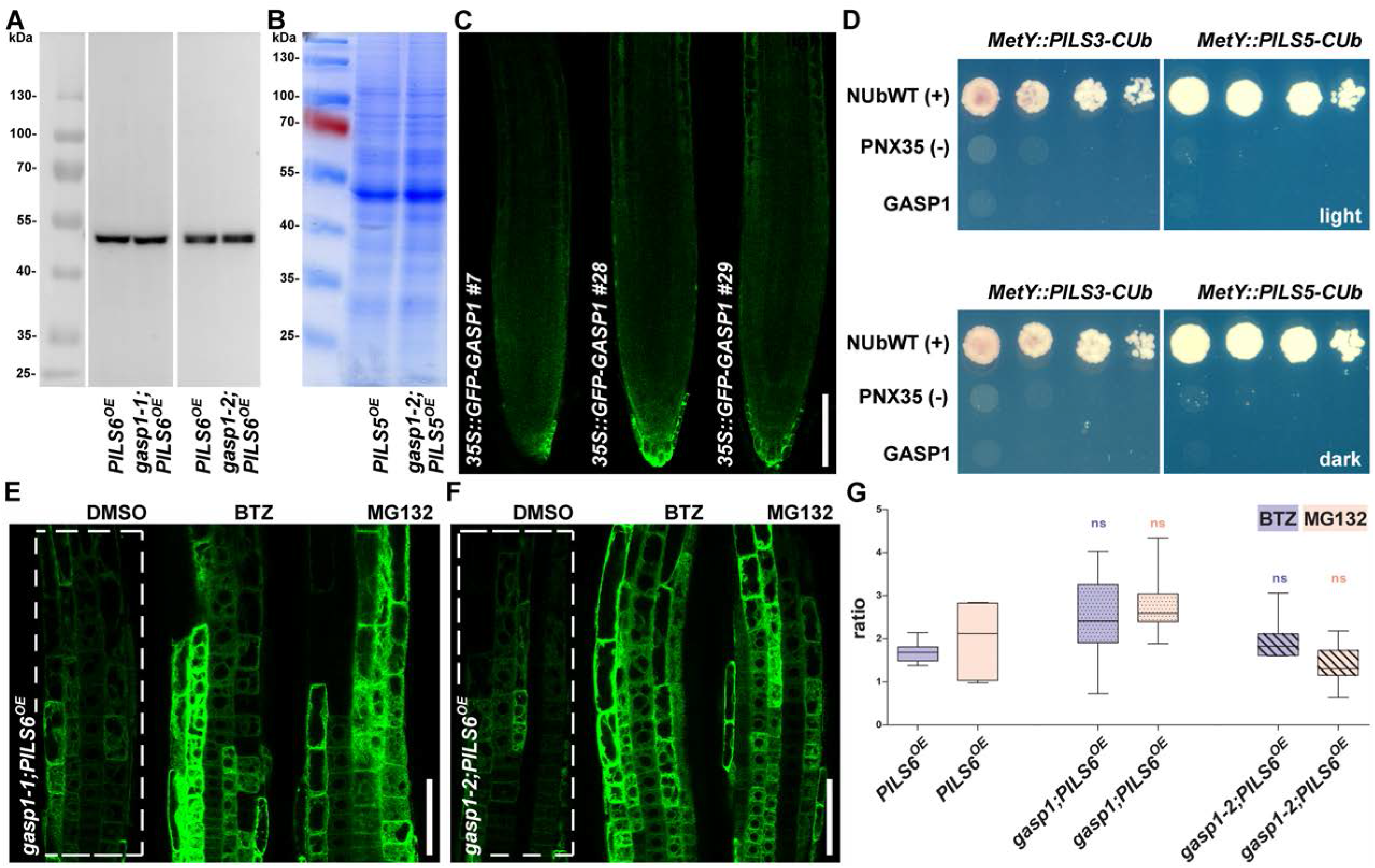
GASP1 affects indirectly the proteasome-dependent PILS6 protein abundance. A,B.Controls for PILS5- and PILS6-GFP Western blots. Anti-Tubulin- (A) and Coomassie- (B) based normalizations were used for the Western blot analyses presented in Figure 3A. C.*35S::GFP-GASP1* localization in roots. Three independent lines show weak but ubiquitous localization in 5 DAG light-grown seedlings. We used mainly line 28. Scale bar, 100 μm. D.PILS3 and PILS5 proteins do not interact with GASP1. Neither PILS3 nor PILS5 interact with GASP1 in the light (upper image) or dark (lower image) in the yeast mating-based split-ubiquitin system. NUbWT was used as a positive control, PNX35 as a negative control. E-G.Proteasome inhibitors stabilize PILS6-GFP independently of GASP1. Confocal images (E, F) and BTZ/DMSO and MG132/DMSO ratios of signal intensity (G) show that a short treatment (3 h) with the proteasome inhibitors Bortezomib (BTZ; [50 uM]) or MG132 [50 uM] stabilizes PILS6-GFP in WT and *gasp1* mutants. The ratios were calculated with the values from Figure 3E. ns = not significant, One-way ANOVA and Tukey’s multiple comparison test (G). Scale bars, 50 μm (E, F). The white, dashed rectangles show the ROIs used to quantify the signal intensity.

**Figure 4S.**
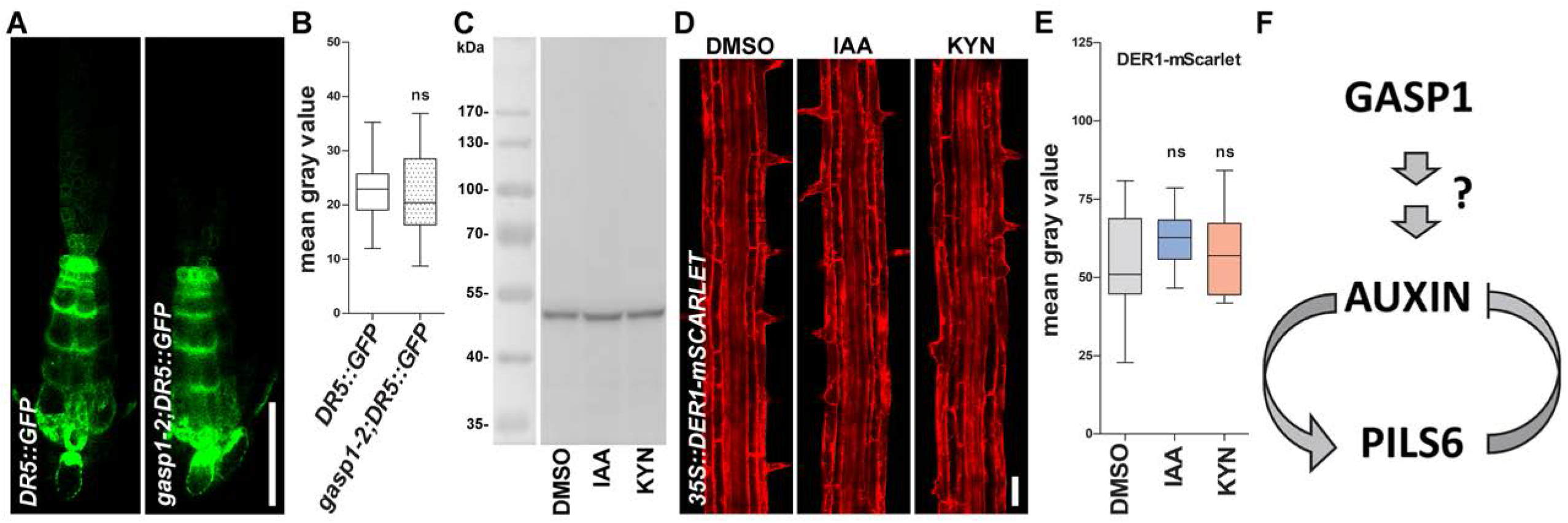
Auxin feedback on PILS proteins. A,B.*DR5::GFP* signal intensity is not affected in the very root tip of *gasp1-2* seedlings. Confocal images (A) and quantification of signal intensity (B) show slightly but not significantly weaker *DR5::GFP* signal intensity in the root tip of *gasp1-2* mutant grown in the light for 5 DAG. n = 15, 16; ns = not significant, t-test and Mann-Whitney test (B). Scale bar, 50 μm (A). C.Anti-Tubulin-based normalization was used for the Western blot analysis presented in Figure 4I. D,E.Auxin signaling does not affect *35S::DER1-mScarlet*. Confocal images (D) and quantification of signal intensity (E) show that a 24 h treatment with either [100 nM] IAA or [1uM] KYN does not affect the fluorescence of DER1-mScarlet. n = 11; ns = not significant, One-way ANOVA and Tukey’s multiple comparison test (E). Scale bar, 50 μm (D). F.Working model illustrating our findings. GASP1 modulates (directly or not) auxin signaling output, which further influences PILS6 protein stability. In return, PILS6 activity represses the abundance of auxin for nuclear auxin signaling. This intracellular feedback regulation between auxin and PILS6 may allow an optimal auxin concentration for fine-tuning auxin-dependent plant responses.

